# Length of Mucin-Like Domains Enhance Cell-Ebola Virus Adhesion by Increasing Binding Probability

**DOI:** 10.1101/2020.07.02.184580

**Authors:** X. Cui, N. Lapinski, X. Zhang, A. Jagota

## Abstract

The Ebola virus (EBOV) hijacks normal physiological processes by apoptotic mimicry in order to be taken up by the cell it infects. The initial adhesion of the virus to the cell is based on the interaction between T-cell immunoglobulin and mucin domain protein, TIM, on the cell-surface and phosphatidylserine (PS) on the viral outer surface. Therefore, it is important to understand the interaction between EBOV/PS and TIM, with selective blocking of the interaction as a potential therapy. Recent experimental studies have shown that for TIM-dependent EBOV entry, a Mucin-like Domain (MLD) with a length of at least 120 amino acids is required, possibly due to the increase of area of the PS-coated surface sampled. We examine this hypothesis by modeling the process of TIM-PS adhesion using a coarse-grained molecular model. We find that the strength of bound PS−TIM pairs is essentially independent of TIM length. TIMs with longer MLDs have higher average binding strengths because of an increase in the probability of binding between EBOV and TIM proteins. Similarly, we find that for larger persistence length (less flexible) the average binding force decreases, again because of a reduction in the probability of binding.

**Statement of Significance:** This work studies the mechanism of TIM-dependent adhesion of the Ebola virus to a cell. Through coarse grained modeling we show that longer TIM stalks adhere more easily as they can sample a larger area, thus offering a mechanistic interpretation of an experimental finding. Better mechanistic understanding can lead to therapeutic ideas for blocking adhesion.

## 1. Introduction

A critical step in the life cycle of a viral particle is its adhesion to and uptake by the cell it infects (1, 2). A virus usually accomplishes this by hijacking normal physiological processes such as endocytosis, phagocytosis, and macropinocytosis (3-6). Ebola virus (EBOV), one of the filoviruses, is a single strand, negative-sense RNA virus that can infect a wide range of mammalian cells. Ebola virus disease is a severe and often fatal illness that can cause hemorrhagic fever in humans. This disease was first identified in 1976, with a fatality rate of 50% to 70%, and has caused over 15000 deaths (7-10). Recent studies show that a set of cellular proteins and molecular mechanisms are required for the virus to enter host cells (6). The internalization of EBOV is initiated by the interaction between viral surface phosphatidylserine (PS) and host cell receptors (11, 12). One type of these receptors are T-cell immunoglobulin mucin domain (TIM) family proteins, TIM-1 and TIM-4 (but not TIM-3). Specifically, attachment involves interaction between TIM proteins and phosphatidylserine molecules displayed on the viral surface (Fig. 1a) (1, 2, 13-15). Once this binding occurs, the virus is internalized into endosomes mainly via macropinocytosis. Therefore, blocking the binding of EBOV to its host cell could be a promising therapy (16). However, a principal difficulty for designing therapies against viruses lies in the fact that any attempt to block the binding of a virus to a cell surface is likely also to deleteriously affect normal physiological processes, such as the TIM-dependent engulfment of dead cell debris by phagocytes. A better understanding of how physical parameters, such as stiffness, length and ligand density, influence the receptor-mediated virus uptake could potentially aid in the development of better antiviral strategies (11, 16).

**Figure 1.**
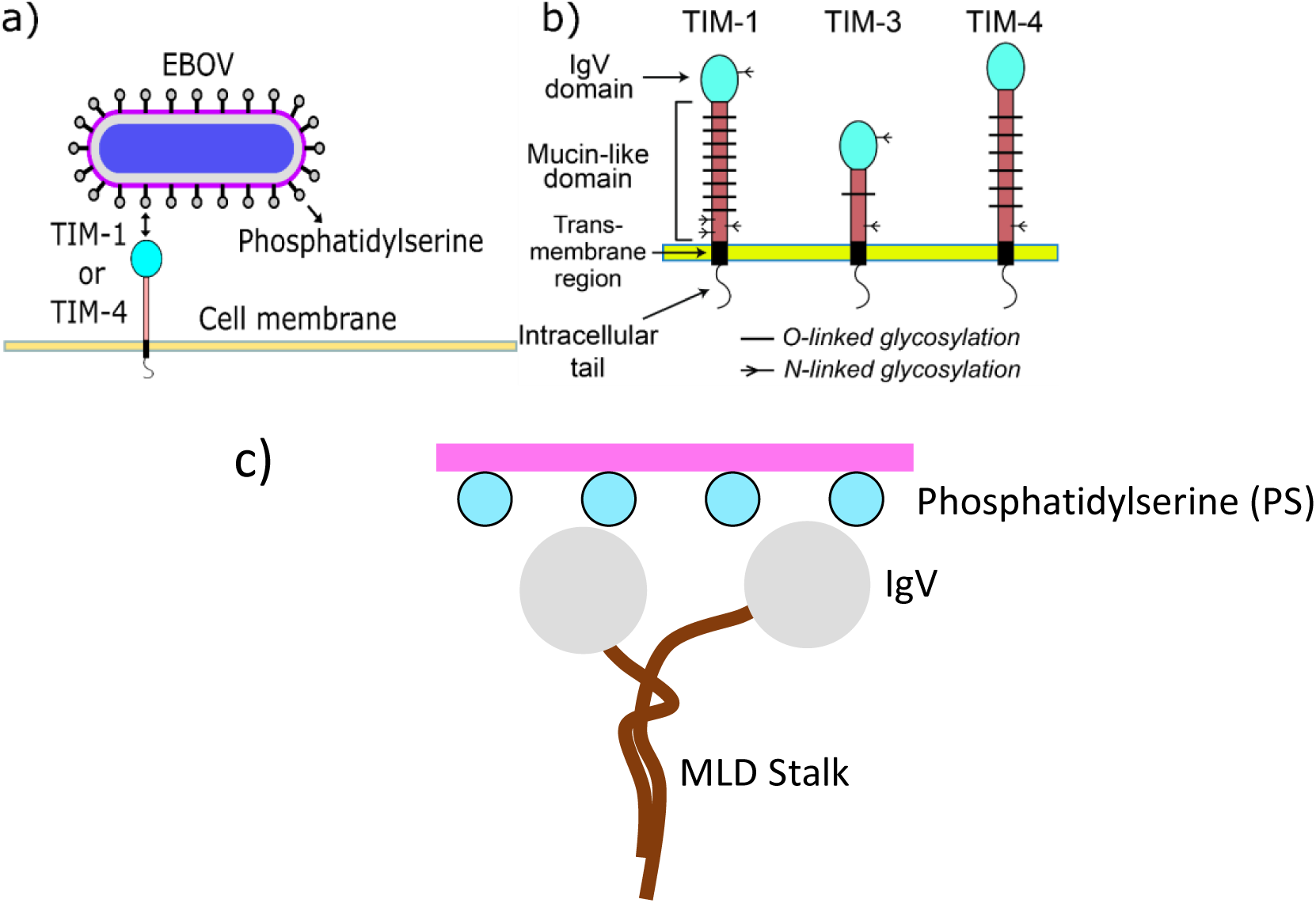
(a) Simplified model for initiation of EBOV uptake (4, 5, 9). EBOV encounters potential host cells and attaches to TIM-1 or TIM-4 on the cell surface via phosphatidylserine. (b) Human TIM protein family. The extracellular portion of human TIM is highly conserved, with an N-terminal IgV domain adjacent to a mucin-like domain (MLD). The IgV domain contains a PS-binding metal-ion-dependent ligand-binding site (MILIBS). The MLDs of TIM-1 and -4 are highly glycosylated (57 and 41 predicted O-glycosylation sites, respectively), resulting in a bulky, unstructured and negatively charged macro glycopeptide region. TIM-2 (not shown) is found only in mice. (c) Hypothesis to explain the advantage of longer MLD: they are more flexibility than shorter MLD proteins, which means they have a larger area of interaction.

Human TIM proteins (Fig. 1b) are a family of type 1 transmembrane receptors widely expressed on cells in the immune system, including T-and B-lymphocytes, dendritic cells and macrophages (17). The IgV and mucin-like domain (MLD, the stalk presenting the ligand-binding IgV domain) comprise the TIM ectodomain and serve essential functions in EBOV uptake. The role of the MLD is poorly understood. The size of the MLD varies among the different TIM family members, with TIM-3 having a short MLD (66 amino acids (AAs)) compared to the longer TIM-1 (167 AAs) and TIM-4 (174 AAs). Recent studies show that MLD length plays a role in TIM-PS binding, and that MLDs of at least 120 AAs are required for sufficient EBOV uptake (16).

One hypotheses to explain the observed effect of MLD length on TIM-PS adhesion is that with increased length comes increased flexibility in the MLD stalk, effectively increasing the region of the viral surface that can be explored by IgV, possibly increasing the likelihood of forming adhesive bonds (Fig. 1c) (12). In this work, we examine this hypothesis by developing a coarse-grained computational model of a single TIM stalk interacting with a plane of PS beads. We simulate the process of binding and separation of the two, as a function of three parameters: MLD length, MLD flexibility, and PS density. In all cases, we find that the adhesion strength of a single IgV-PS interaction, as measured by the force required to separate IgV from PS, is essentially unchanged if preceded by a binding event. The probability of binding itself depends strongly on each of the three parameters, with the net effect that increase in average separation force with increasing MLD length is essentially entirely due to a larger probability of binding to a PS bead.

## 2. Methods

We simulated the process of adhesion and separation by first constructing a coarse-grained model of the TIM-PS system (Fig. 2). Our model captures the essential physical features: (a) a plane embedded with PS molecules located a specified distance apart, represented by a single bead each, (b) the terminal IgV domain of the TIM protein, represented by a large bead, and (c) the stalk-like MLD represented by a string of connected beads.

**Figure 2.**
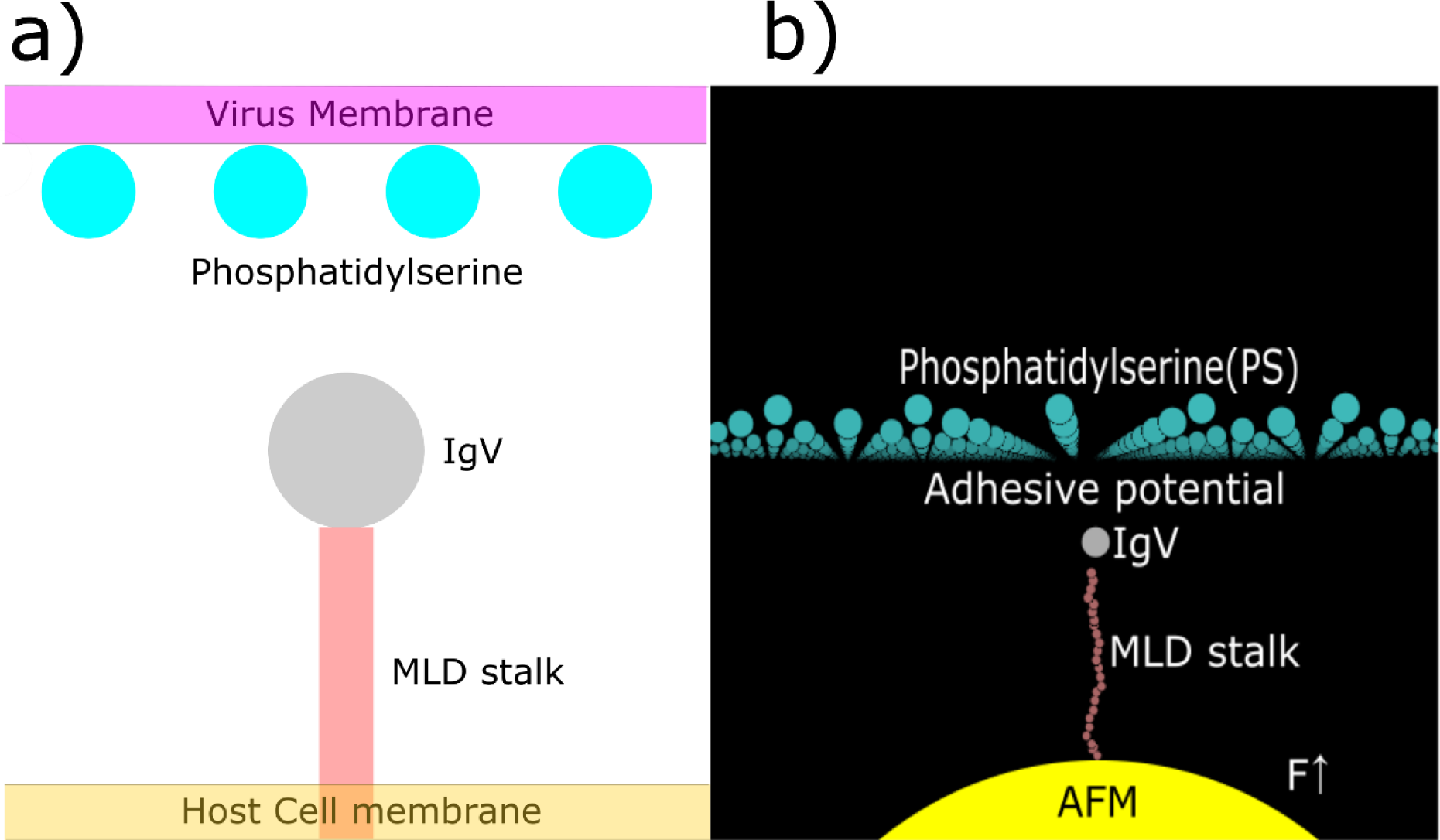
Coarse-grained model that captures the main physical features of TIM-PS interaction. (a) The essential features are shown schematically. They comprise a host membrane in which the TIM receptor is embedded, which binds to phosphatidylserine on the virus surface via the IgV bead. (b) Coarse-grained representation comprising (i) a single IgV bead, (ii) a string of MLD beads, (iii) a single bead representing AFM force probe subjected to force *F*, and (iv) an array of PS beads embedded in a plane representing the viral surface.

Multiple potentials were applied to the system, as summarized in Table 1. The adhesive interaction between IgV and PS was represented by a Born-Mayer-Huggins function with a potential depth of 50 k_B_T (300 K) and minimum of potential at 3.5 nm (equilibrium distance of separation without external load (18)). The MLD was represented by a string of connected beads (R_MLD_=0.5 nm) with stiff spring potentials between adjacent beads and bending potential for each set of three adjacent beads. The spring constant *K*_*s*_ was set at a high value (100 N/m) to constrain the inter-bead distances to be near their natural length L_0_ (19). The bending constant *k*_*θ*_ was in a range set by the MLD stalk’s known persistence length (19-21) and the minimum of the bending potential was at an angle *θ*_0_ of 180 degrees, i.e., for a collinear configuration of the three beads. A background long-range repulsion from the PS plane against all the beads was used to represent repulsion from glycoproteins on the viral surface between PS beads that prevent any of the MLD beads from penetrating the virus membrane. One additional bead with large radius (R=50 nm >> R_IgV_=3.5 nm) was placed above the PS plane to ensure that the IgV bead does not cross the membrane. We additionally included repulsion between MLD-MLD and MLD-IgV beads to establish self-avoidance. A single simulation began with the TIM beads being pushed towards the PS plane and ended with them being pulled away. This was achieved by attaching the lowest MLD bead to a large bead (e.g., representing an AFM tip (R_AFM_=40 nm)), mimicking a single-molecule AFM experiment (20, 22-24). A time-dependent force was applied to this AFM bead to drive the process of approach and retraction of TIM from the PS plane.

**Table 1:**
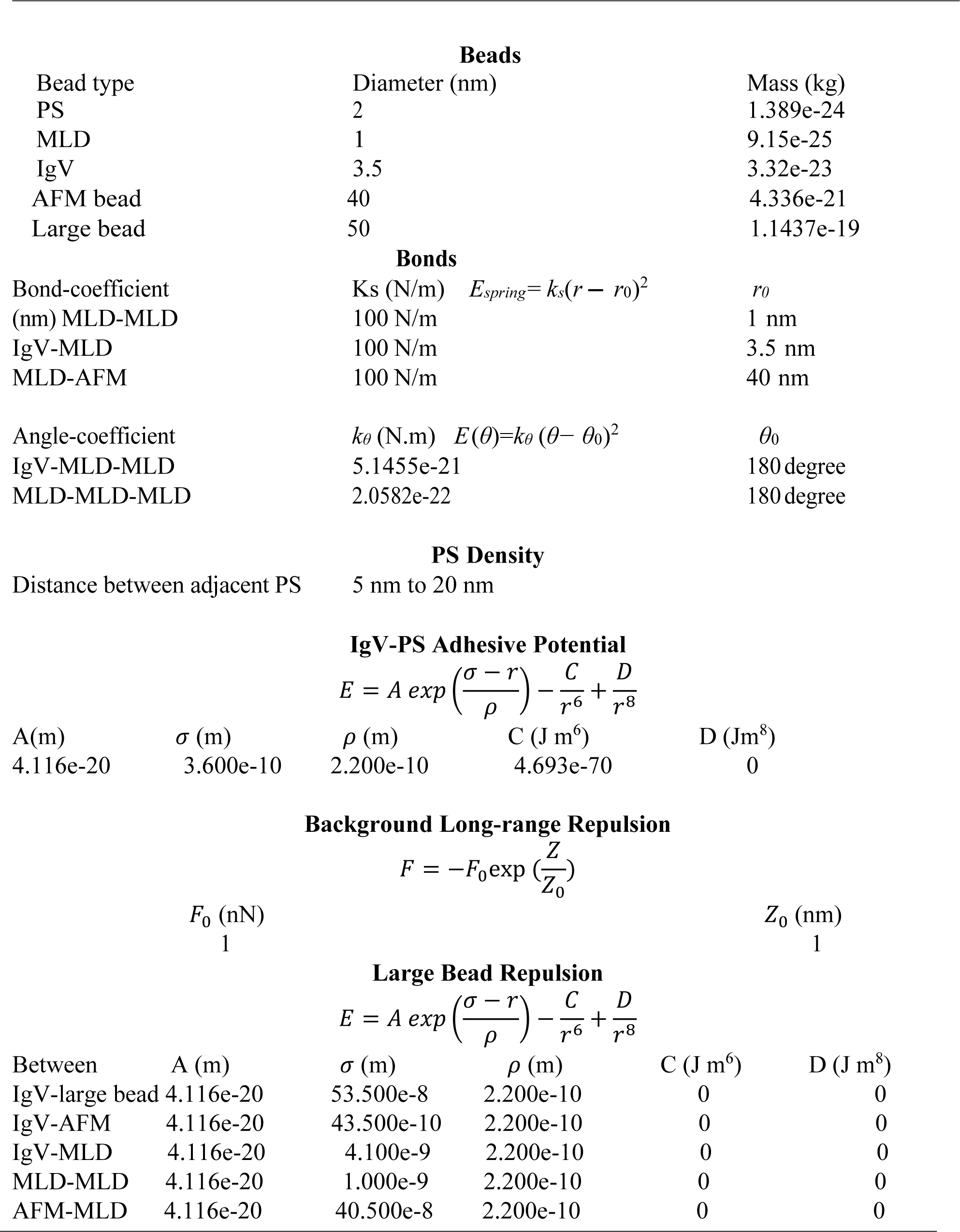
Parameters for Potentials and Forces in the Coarse-Grained Model

Using the model just described, we simulated the time-dependent process in which the TIM receptor is driven by the AFM bead towards the PS surface and then retracted (as might occur in a single-molecule AFM experiment) (20). The adhesive potential depth was adjusted to 50 k_B_T to bring the computed pull-off forces within an experimentally relevant range at the loading rate used in the simulation. The model was implemented and simulated through LAMMPS (Large-scale Atomic/Molecular Massively Parallel Simulator) (25, 26) and the results were analyzed using scripts written in MATLAB and VMD (Visual Molecular Dynamics) (27).

A time-dependent force over a range of loading rates was applied to establish a range of pull-off forces. Then, for the majority of simulations in which parameters were varied, we used a loading rate of 0.0003 N/s for 170 ns and then -0.0003 N/S for 830 ns. The former positive loading rate represents pushing the TIM molecule towards the PS plane while the latter represents pulling the molecule away.

Fig. 3 shows six snapshots from a typical simulation. Fig. 3a (t=0 ns), represents the initial state in which the MLD beads are arranged in a straight line and the ramp force has just begun to be applied to the large yellow (AFM) bead. At t=100 ns, the force applied to the AFM bead has pushed up the lower end of the MLD chain. At t=150 ns, the MLD chain has been pushed against the PS-embedded surface. In this case (but not in all) the IgV bead binds to one of the PS beads on the viral surface. At t=450 ns, the AFM bead is now subjected to a pulling force. The MLD has started to be pulled with the IgV bead still bound to the PS bead. At t=600 ns, with continually increasing pulling force, the MLD length starts to be stretched but the IgV bead is still bound to a PS bead. At t=800 ns the unbinding process has completed, and the IgV-MLD chain is pulled away from the PS plane. In SI we provide two videos, one representing the cases where the IgV bead does and a second one for cases where it does not bind to a PS bead during this process.

**Figure 3.**
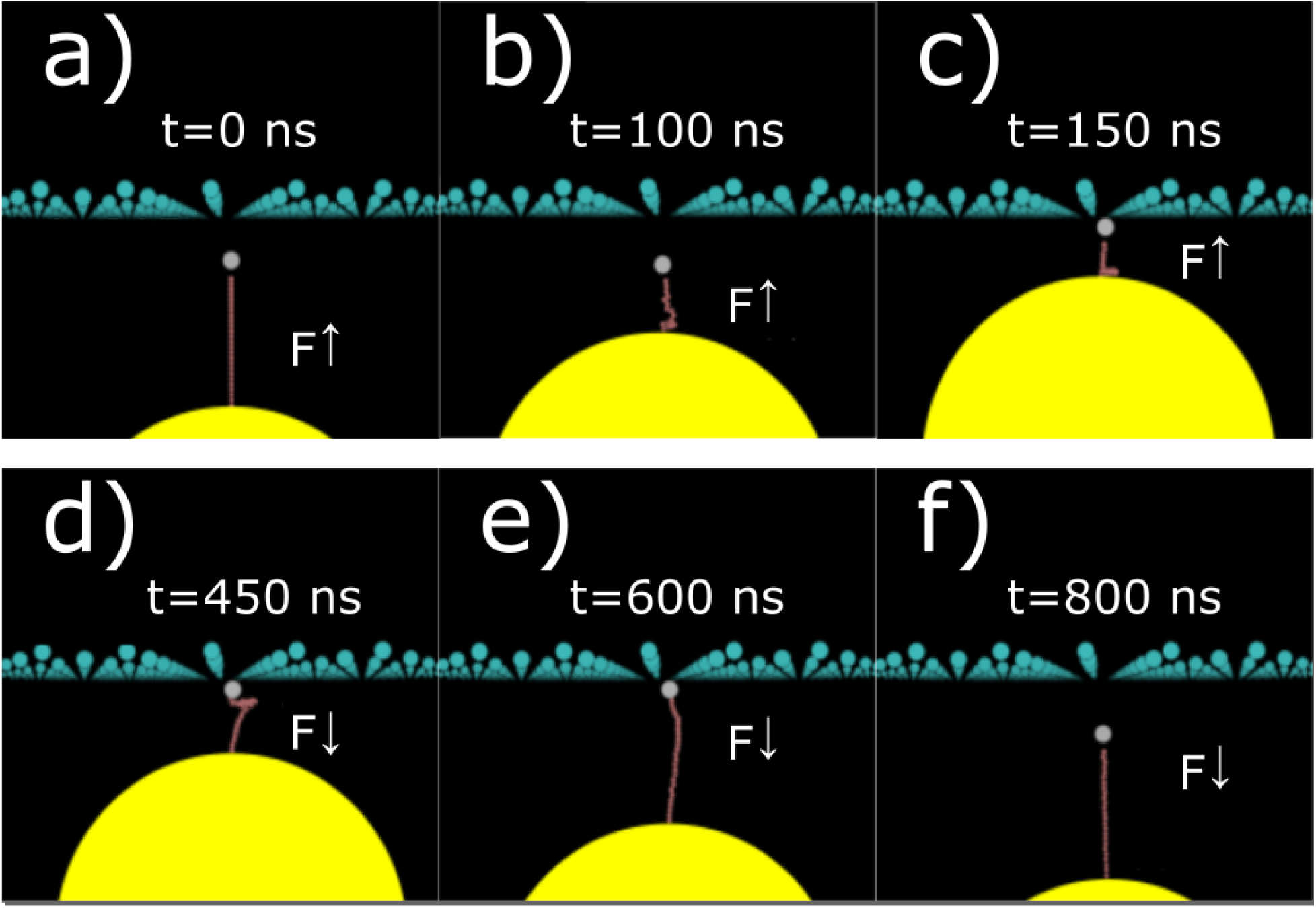
Snapshots from different stages of a typical simulation. a) The initial state in which the MLD beads are arranged in a straight line and the ramp force has just begun to be applied. b) The force applied to the AFM bead has pushed the end of the MLD chain and conformations of the MLD begins to randomize. c) The MLD chain is pushed against the PS-embedded surface. d) The MLD has started to be pulled with the IgV bead still bound to a PS bead. e) The MLD chain begins to be stretched while the IgV bead is still bound to a PS bead. f) The unbinding process is completed, and the IgV-MLD chain is pulled away from PS plane. Arrows indicate the direction of force applied to the AFM bead.

Before conducting production simulations, we needed (a) to establish a loading rate that resulted in appropriate values for the pull-off force, (b) to confirm that the pulling rate was sufficiently slow so the pull-off force was governed by the kinetics of hopping over the energetic barrier, and not by the drag force, and (c) to establish a criterion by which we determined whether the IgV bead did or did not bind to a PS bead prior to being pulled away. In Fig. 4, we plot the distance of several beads from the PS plane as a function of pulling time. The number in the legend represents the bead number such that ‘1’ corresponds to the IgV bead and 50 is the bottom MLD bead attached to the AFM bead. We found that at this chosen rate (0.0005 N/s) the distance versus time plot for bead ‘1’ remained relatively unchanged until about 500 ns of pulling, which corresponded to the IgV bead being bound to a PS bead for some time before dissociating from it. In contrast, the bottom MLD bead was pulled down immediately when the pulling force is applied, but it also had an intermediate plateau during which period the molecule was stretched but remained bound. This showed that the pulling rate was slow enough that the MLD chain had sufficient time to sample configurations in the bound state before being pulled off.

**Figure 4.**
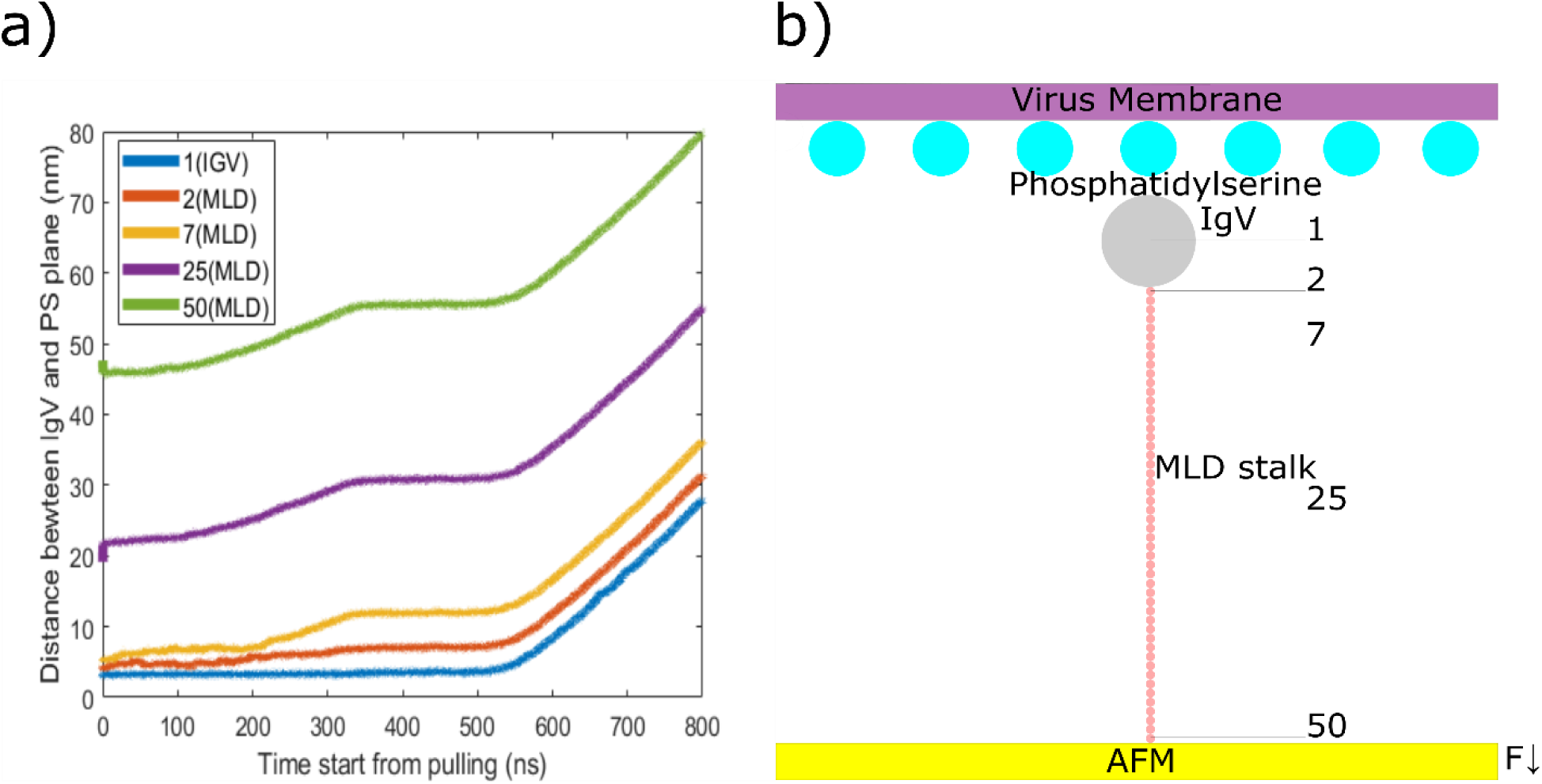
(a) Demonstration that unbinding is governed by hopping over the energetic barrier rather than drag force: The bottom MLD bead (#50) starts to move once pulling force is applied while the top MLD bead (#2) and IgV (#1) move little until the pulling force reaches a sufficient value, which indicates that IgV is captured by the energetic barrier. (b) Schematic of unbinding system corresponded to data in (a).

Next, we needed to develop a method to calculate the pull-off force and to set up a criterion to distinguish between cases in which unbinding was preceded by an IgV-PS binding event from those in which it was not. The pull-off force was taken to correspond to the time point at which the z-position of the IgV bead relative to the PS plane became sufficiently large. Specifically, we defined separation as when the IgV-PS-plane distance exceeded twice the equilibrium value of 3.5 nm. Choosing this number between 6 and 10 nm resulted in no qualitative differences and only small quantitative differences in our results. If *t*\* was the time at which this occurred, the corresponding pull-off force was calculated as:

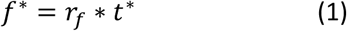

where *f*^*\**^ is the pull-off force, *r*_*f*_ is the constant loading rate and *t*^*\**^ is the time from the beginning of pulling to the time when unbinding occurred.

Whether the IgV bead was able to find a PS binding partner prior to being pulled away was easily observed by viewing a video of the simulation. (See Supporting Information for two videos, one in which the IgV bead finds a PS binding partner and one in which it does not.) To determine from the pull-off force whether it corresponded to the removal of a PS-bound or PS-unbound IgV bead, we show in Fig.5 the distribution of the pull-off force.

**Figure 5.**
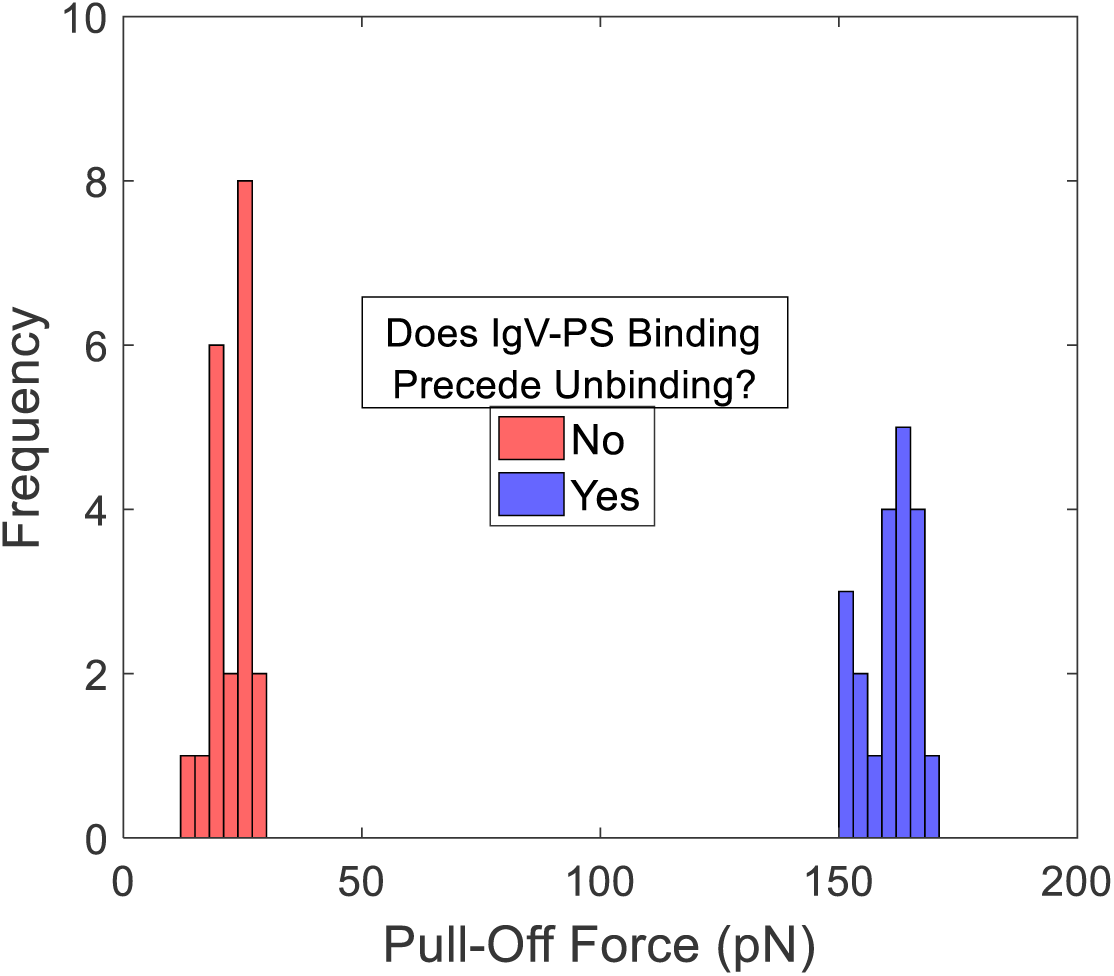
Criterion to determine if unbinding was preceded by an IgV-PS binding event. Pull-off forces clearly falls into two distinct and separate classes. When unbinding is preceded by an IgV-PS binding event, pull-off forces are always >120 pN. Conversely, when unbinding is not preceded by IgV-PS binding, pull-off forces are always <30 pN. Therefore, a threshold value of pull-off force between 30 and 120 pN can safely be used to classify each simulation according to whether or not unbinding is preceded by an IgV-PS binding event.

It is evident that the distribution is strongly bimodal. The pull-off forces in all the cases where IgV was not bound to a PS bead prior to unbinding were below 30 pN. In all the cases where the two were bound prior to unbinding, the pull-off forces were greater than 120 pN. Therefore, we could safely identify all the latter cases as those with pull-off force > 80 pN and the former cases as those with pull-off force < 80 pN.

Finally, we conducted a series of simulations at different pulling rates using a single IgV bead to investigate the rate dependence of pull-off force and to choose a loading rate for the rest of our study. We used 120 AA MLD as a test example. These simulations began with the IgV-MLD situated under a PS bead at a distance of 3.5 nm. The pushing phase was kept at the loading rate of 0.0005 N/S for 200 ns but we varied the pulling rate from -0.0005 N/s, to -0.01 N/s, repeating each case 5 times. The average pull-off force is plotted against the logarithm of the pulling rate in Fig. 6.

**Figure 6.**
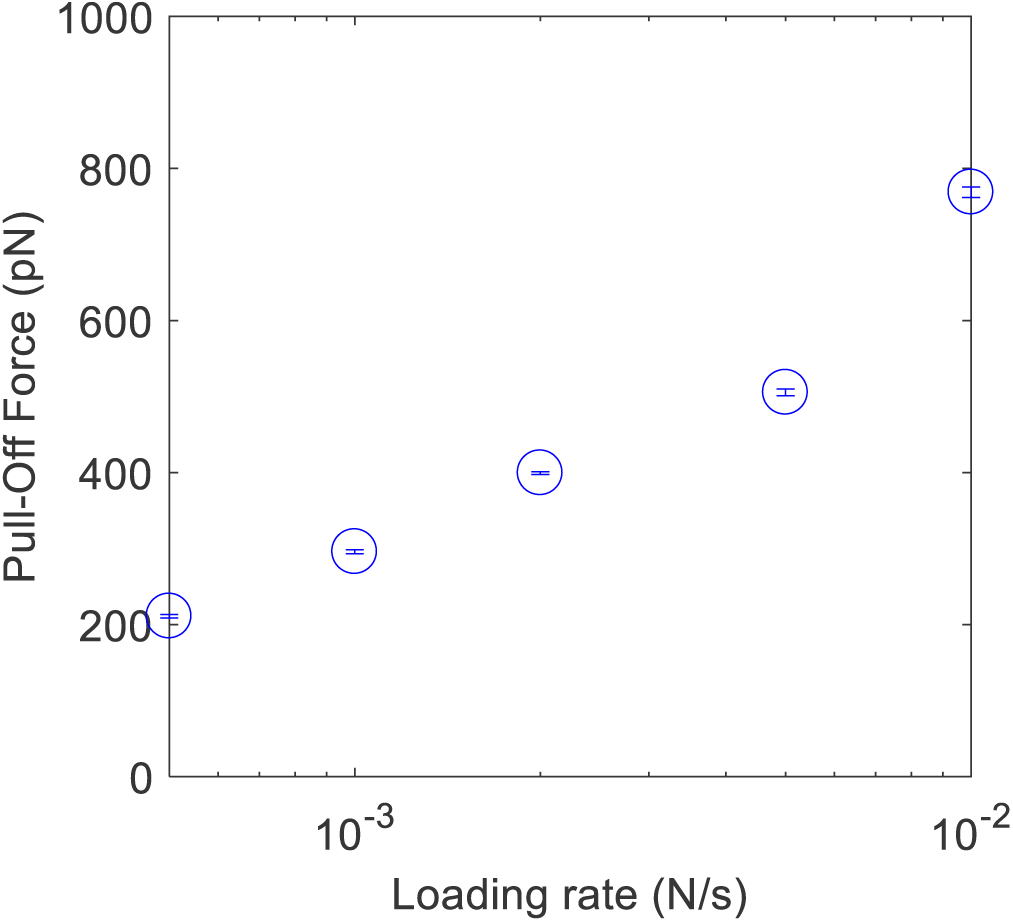
Average pull-off forces for different loading rates are plotted against the corresponded loading rates. The plot shows that pull-off force is approximately linear in logarithm of loading rate.

These results are consistent with the Bell-Evans model with constant loading rate, which predicts that (20):

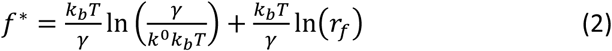

where *k*_*b*_ is Boltzmann constant, T is the absolute temperature, *k*^0^ is the dissociation rate constant without a pulling force and γ is the position of the transition state. As predicted by this model, the un-binding force *f*^*\**^ is linear in the logarithm of the loading rate. Moreover, at the lowest loading rate, the pull-off forces were within the range obtained by single-molecule experiments.

The important parameters relevant to the hypothesis we explored in this work that can be varied in our coarse-grained model are the MLD length, MLD persistence length, and PS density. We first explored the effect of the MLD length on the pull-off force, holding the PS density and persistence length constant, by varying the MLD length from 5 to a maximum of 250 AAs. In addition to varying the MLD properties, we also randomly sampled the X-Y location of the lowest MLD bead. We conducted 50 simulations for each MLD length. To study the effect of MLD persistence length, we fixed the PS density and MLD length (120 AAs), but varied the MLD persistence length from 0.01 nm to 100 nm. Similarly, the effect of PS density was explored by varying the distance between adjacent PS beads with 40 simulations for each PS density. Each case was repeated 35 times.

## 3. Results and Discussion

### 3.1. Effect of MLD Length

Fig. 7 shows the results of simulations in which the MLD length was varied. We calculated the average pull-off force in two ways, (a) average over all the simulations, including IgV-PS bound and unbound cases, and (b) average over only those cases where pull-off was preceded by successful binding of IgV to PS. We also present the probability of IgV-PS binding for each MLD length.

**Figure 7.**
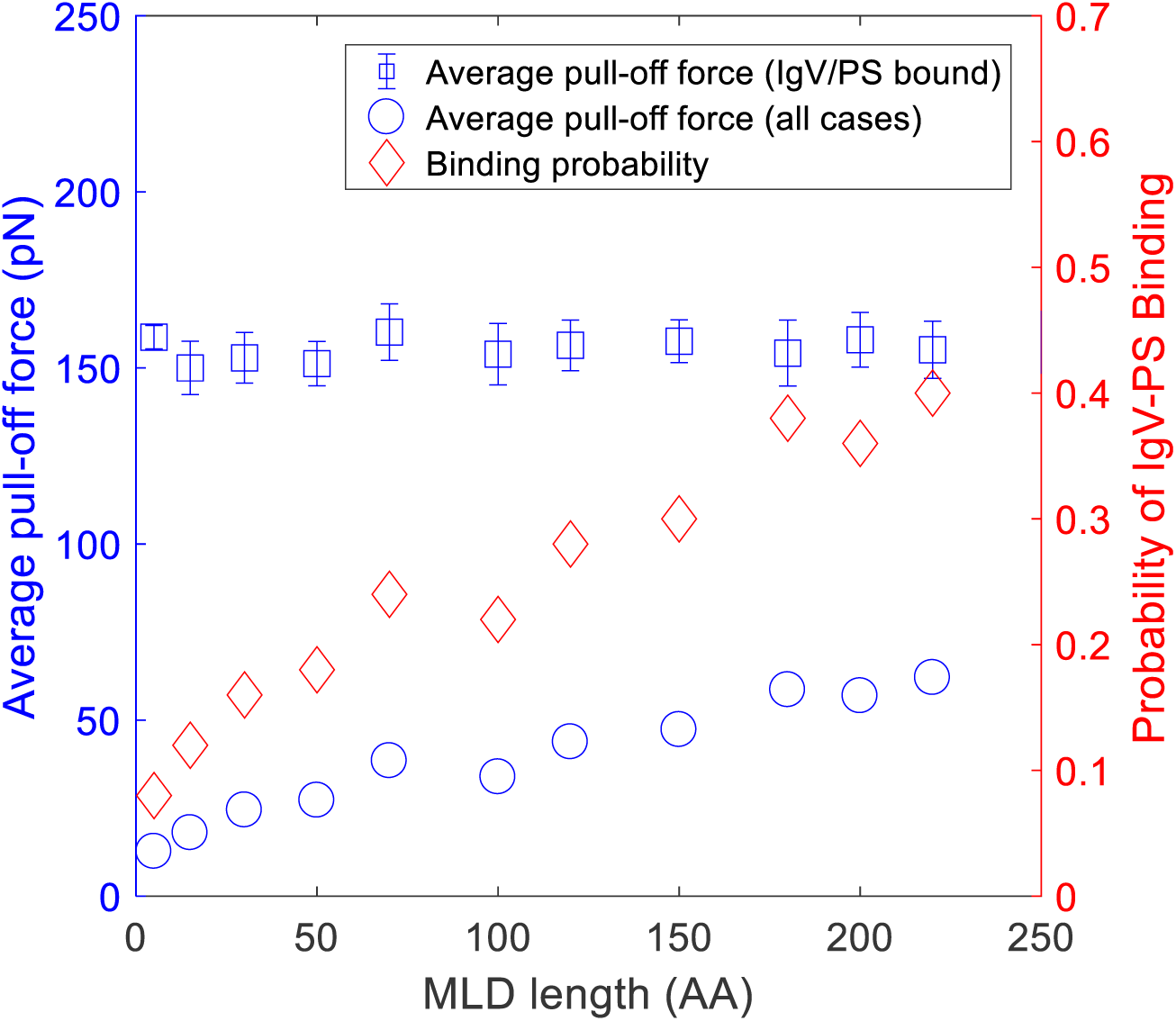
The average pull-off force for all cases (blue circles) increases with longer MLD. In contrast, the average pull-off force (blue squares) for only those cases where IgV-PS are bound prior to being pulled-apart has roughly a fixed value. This indicates that the increase in pull-off force for longer MLDs is essentially entirely due to a higher probability of IgV-PS binding (red rhombuses). This supports the hypothesis that increasing MLD length helps to sample and find PS partners for IgV.

The binding probability increases systematically and significantly with increasing MLD length. This is in agreement with, and supports the hypothesis that, a longer MLD length allows sampling of a larger area on the viral surface, resulting in greater probability of binding. The effect of this on the average pull-off force is striking. The pull-off force, averaged over all simulations, follows the trend set by the probability of IgV-PS binding. In contrast, the pull-off force averaged over only those cases where IgV-PS binding preceded pull-off shows that the strength of binding, once IgV does bind to PS, is essentially independent of MLD length. This is far from obvious because with increasing length the MLD stalk does sample and attach to PS beads over an increasing area. However, during pulling, the surface adsorbed portion of the MLD chain slides so that just prior to attachment the IgV bead is generally directly above the pulling bead. Thus, the process begins with stochastic uncertainty about whether IgV binds to PS. If it does not bind, IgV is pulled off at a very low force. If it does bind, it takes a much larger pull-off force and in the process, IgV loses its memory of where it first attached. Thus, we can say succinctly that average *F*_*pull*−*off*_ = *F*_*pull*−*off*_^*\**^ *\* Prob*(*IgV* − *PS binding*). The first term on the right-hand side, *F*_*pull*−*off*_^*\**^, is a property of the binding pair, unaffected by MLD length.

### 3.2. Effect of MLD Persistence Length

Fig. 8 shows the results for the effect of MLD persistence length on the binding probability and pull-off force. We found that the binding probability decreases systematically with longer MLD persistence length. Evidently, similar to the effect of MLD length, shorter persistence length means that the more flexible MLD stalk allows IgV to sample a larger area. Precisely as in the case of MLD length, the average pull-off force for cases where IgV-PS binding precedes unbinding is essentially independent of persistence length. Again, this suggests that the flexible MLD chain fosters binding by facilitating IgV access to a larger region.

**Figure 8.**
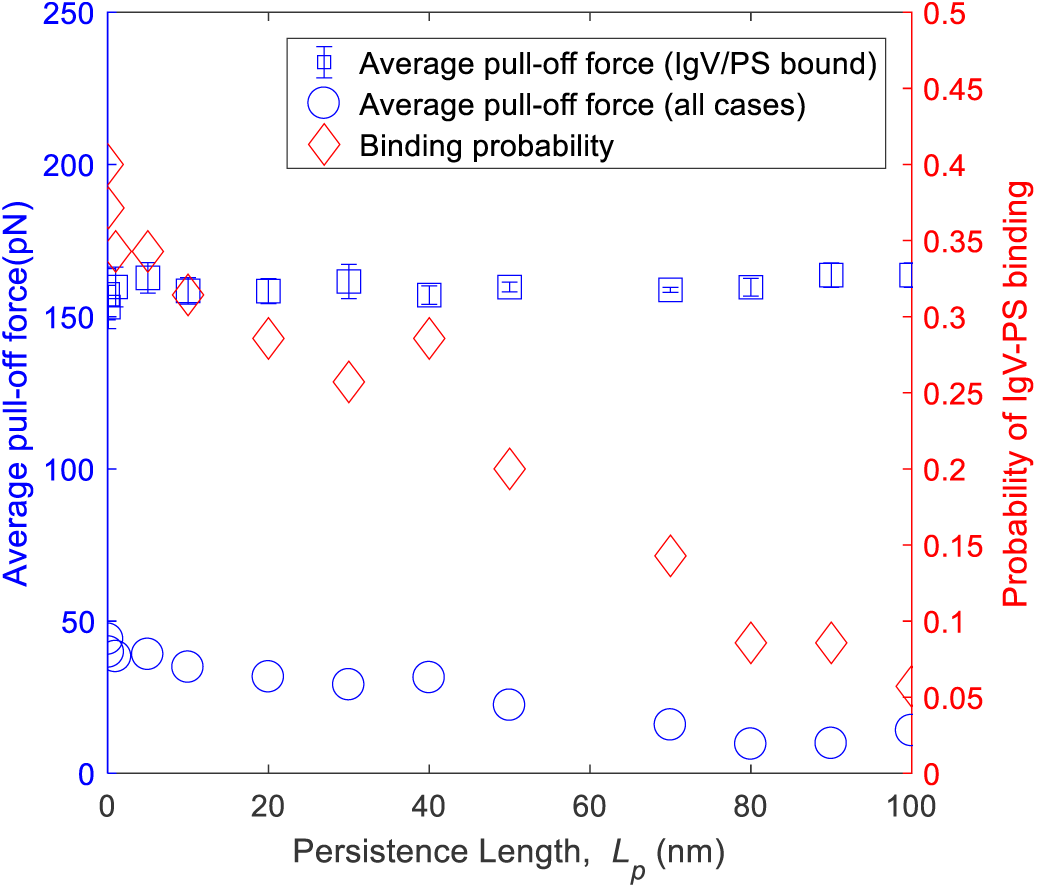
The pull-off force averaged over all simulations (blue circles) decreases with increasing MLD persistence length. However, the pull-off force averaged over only the simulations where the IgV bound to PS (blue squares) remains roughly unchanged, which indicates that the decrease in the former for a higher persistence length is due to a reduction in the binding probability (red rhombuses). That is, a flexible MLD stalk (small persistence length) favors PS-IgV binding.

### 3.3. Effect of PS Density

Fig. 9 shows the effect of PS density on binding probability and pull-off forces. We found that the binding probability decreases with decreasing PS density, due to a larger area that the IgV needs to sample to find a PS binding partner. Consistent with the effects of MLD and persistence lengths, the pull-off force averaged over cases where unbinding is preceded by IgV-PS binding is independent of PS density. Thus, the effect of decrease in PS density operates by increasing the area that IgV needs to sample.

**Figure 9.**
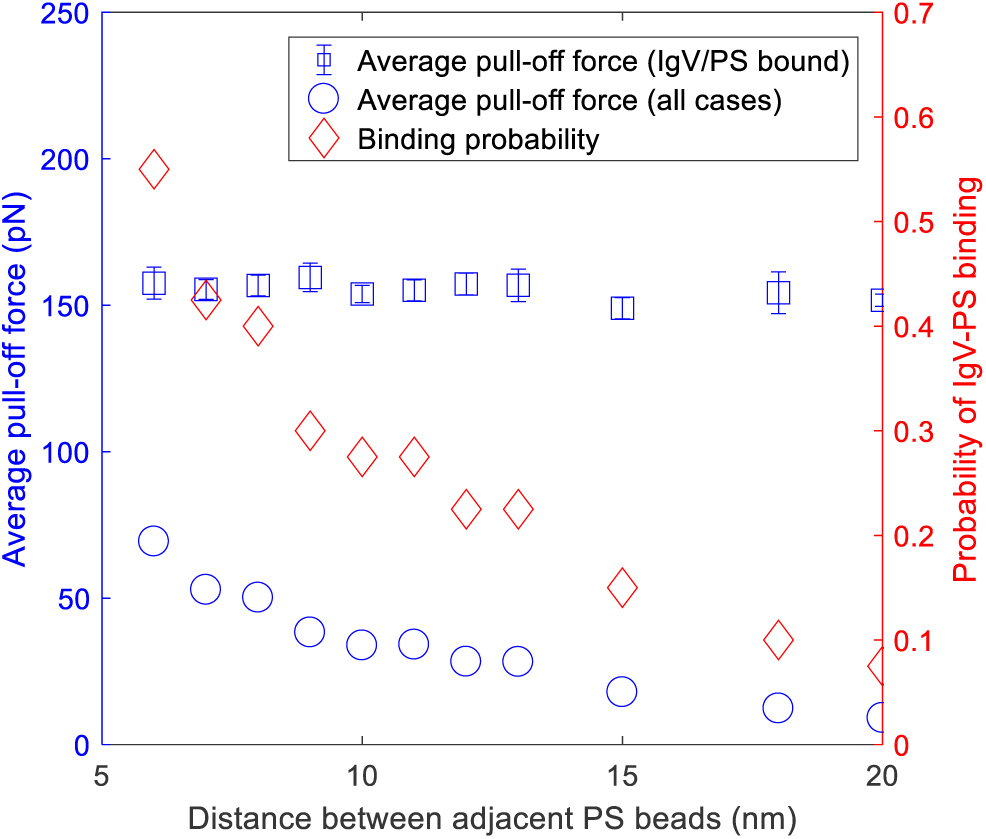
Binding probability and average pull-off force for different PS densities. The plot shows that a higher PS density significantly increases the IgV-PS binding probability, while the average pull-off force for cases where IgV-PS binding occurs remain roughly constant.

## 4. Conclusion

In this study, we built a coarse-grained model for the TIM-PS interactions that enable initial EBOV adhesion to cell surfaces. We used the model to examine a hypothesis that explains the experimental finding that MLD stalks longer than a certain length are required for successful TIM-PS binding. The hypothesis we examined posits that longer MLD stalks bind more effectively because they can sample a larger area on the viral surface. To examine this hypothesis we constructed a model in which the MLD was represented by a connected string of beads, terminated at one end by a larger bead representing IgV. We conducted coarse-grained Brownian dynamics simulations of pushing a single TIM receptor against a plane embedded with PS beads, followed by retraction to pull-off the TIM away from the PS-embedded surface. We studied the effect of MLD length, persistence length of the MLD, and PS density.

We found that increase in MLD length, decrease in MLD persistence length, and increase in PS density all increase the probability of IgV-PS binding. However, the pull-off force, once IgV-PS are bound to each other, remains unchanged. The results are consistent with the experimental finding that longer MLD is needed for successful TIM-PS binding. It also supports the hypothesis that increasing length of MLD is needed to provide flexibility for the TIM stalk to effectively seek binding partners. Smaller persistence lengths and higher PS density have a similar effect on the binding probability.

In this work, we considered an isolated TIM/MLD stalk. In actuality, TIM is embedded in a brush of other sugar-like molecules. These affect the flexibility of the MLD stalk which, to a certain extent, could be captured by modulating the persistence length of the MLD. However, these molecules likely also provide steric and electrostatic repulsion against the PS embedded surface. This is not accounted for in our model and will be examined in future work.

Our modeling approach contains several elements that we believe can be adapted potentially to study other viruses. For example, SARS-CoV-2 binds through spike proteins to cell membrane-bound receptors and the approach we have taken can be adapted to study the mechanisms of viral adhesion to the cell surface in that case. By elucidating the mechanisms of adhesion, we hope to spark ideas for therapeutic targets.

## 5. Author Contributions

XC and NL developed the model and the codes. XC carried out all the simulations and much of the data analysis. AJ and XZ designed the study, mentored and supervised XC and NL, co-interpreted results and co-wrote the manuscript.

## 6. ACKNOWLEDGEMENTS

This work was supported by the NIH grant 1 R15 AI133634-01A1 and by NSF grant 1804117

